# Definitive demonstration by synthesis of genome annotation completeness

**DOI:** 10.1101/455428

**Authors:** Paul R. Jaschke, Gabrielle A Dotson, Kay Hung, Diane Liu, Drew Endy

**Affiliations:** Bioengineering Department, Stanford University, Stanford, CA, USA; Department of Molecular Sciences, Macquarie University, Sydney, NSW, Australia

## Abstract

Bacteriophage øX174 was the first DNA genome to be sequenced. The genome is well studied by classical methods and is known to encode 11 essential genes. At least 23 closely-related *Bullavirinae* genome sequences are now available. We identified 315 potential open reading frames (ORFs) within the genome via bioinformatic analysis, and a subset of 82 highly-conserved ORFs that have no known gene products or functions. Using genome scale design and synthesis we made a mutant genome in which all 11 essential genes are simultaneously disrupted, leaving intact only the 82 conserved-but-cryptic ORFs. The resulting genome is not viable, as expected. Cell-free gene expression followed by mass spectrometry revealed only a single peptide expressed from both the cryptic-ORF and wild-type genomes, suggesting a potential new gene. A second synthetic genome in which 71 conserved cryptic ORFs were simultaneously disrupted is viable but with ~50% reduced fitness relative to the wild type. However, rather than finding any new genes, repeated evolutionary adaptation revealed a single point mutation modulating translation of gene H, a known essential gene, that fully suppressed the fitness defect. Taken together, we conclude that the annotation of ORFs for the øX174 genome is formally complete. Sequencing and bioinformatics followed by synthesis-enabled reverse genomics, proteomics, and evolutionary adaptation can definitely establish the sufficiency and completeness of natural genome annotations.

## Introduction

Ongoing improvements in DNA sequencing tools have allowed researchers to begin to catalog the full diversity of genetic information in nature (Carlson 2003; Landenmark, Forgan, and Cockell 2015). However, assigning biological functions to the so-revealed sequence data has progressed less quickly and surely. For example, genomes are automatically annotated using computational methods that leverage what is known of conserved molecular mechanisms and that incorporate training data from an increasing number of annotated genomes. Nevertheless, accurate prediction of protein-coding open reading frames (ORFs) remains challenging (Tatusova et al. 2013). One particular challenge is that, typically, only a small subset of apparent ORFs actually encode proteins. Moreover, the quantitative scoring of detailed sequence characteristics for ORFs appears as a continuum, from ORFs that do not encode proteins to full-fledged protein-coding ORFs (Carvunis et al. 2012).

Bacteriophage øX174 has been useful as a model system for genetics research, from before the sequencing era through today. The number of identified protein-coding ORFs in øX174 increased as methods were developed and applied. For example, successive forward genetic screens enabled identification of 10 protein-coding ORFs in the absence of genome sequence information (Hutchison 1969;Burgess and Denhardt 1969;Jeng et al. 1970;Mayol and Sinsheimer 1970; Benbow et al. 1971;Linney, Hayashi, and Hayashi 1972). Sequencing of the 5,386 nucleotide øX174 genome (Sanger et al. 1977) and later comparison to the related G4 phage genome enabled identification of one additional protein-coding ORF (Tessman, Tessman, and Pollock 1980). Today, the øX174 genome is still recognized as encoding only these 11 proteins (Fig. 1A) (Fane et al. 2005). In contrast, an unfiltered view of the øX174 genome reveals up to 315 ORFs. While most of these ORFs are presumably not protein-coding (Fig. 1B), it is not absolutely certain that every protein-coding ORF in the øX174 genome has been discovered.

**Figure 1.**
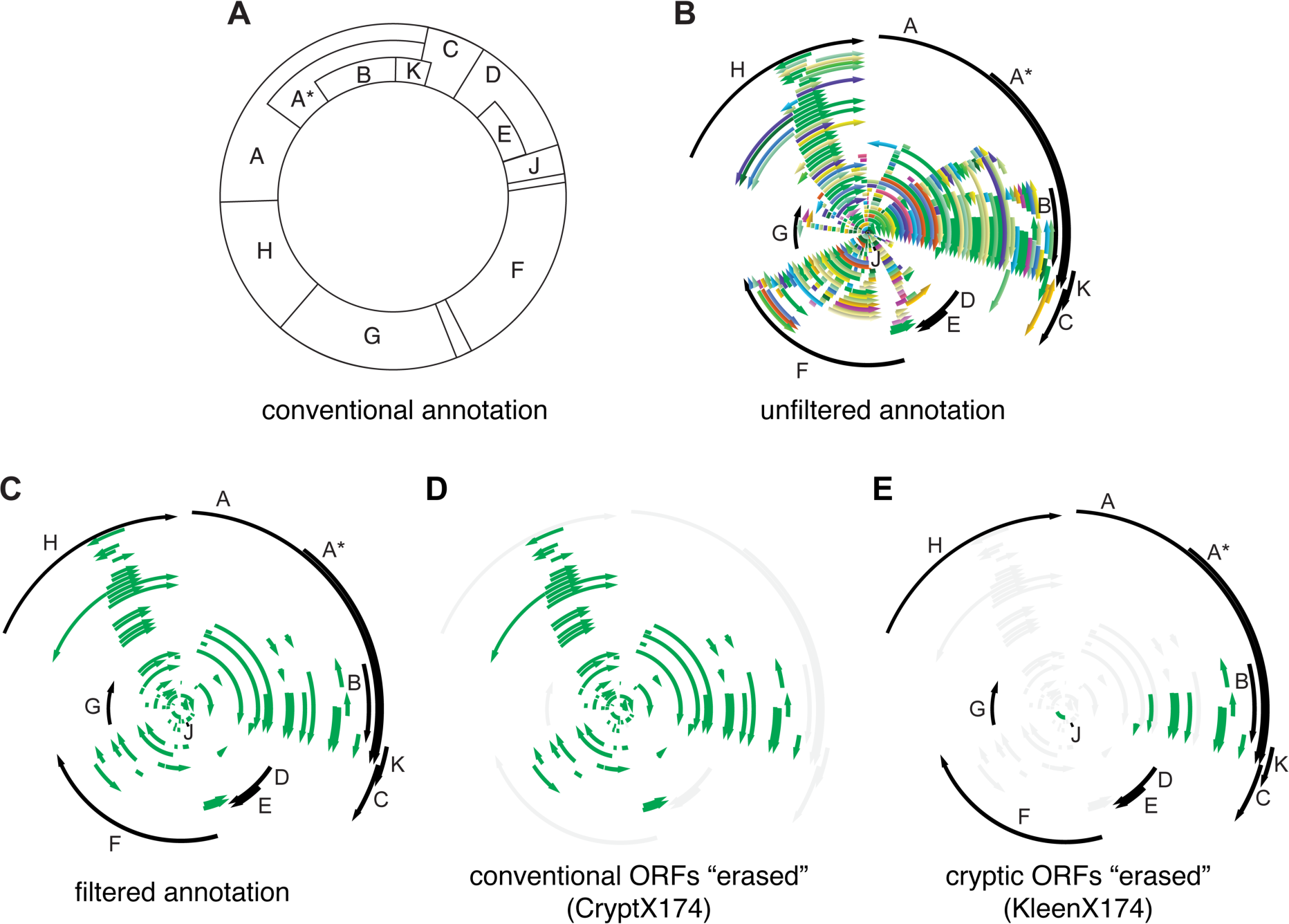
Designing genomes encoding only essential or only cryptic ORFs. (A) contemporary genetic map of øX174. Lettered boxes represent the 11 established protein-coding open reading frames (ORFs). (B) the øX174 genome with 315 ORFs, each 60 base pairs or longer and starting with an ATG, GTG, or TTG codon (various colors), plus the 11 established protein-coding ORFs (black). (C) filtered annotation of the øX174 genome with ORFs; eleven previously identified protein-coding ORFs (black) and 82 cryptic ORFs of unknown protein-coding status (green). (D) cryptX174 genome design with the 11 established ORFs disrupted (gray). (E) kleenX174 genome design showing disruption of 71 cryptic ORFs (gray).

De novo synthesis of the øX174 and other genomes is becoming routine (Smith et al. 2003; Richardson et al. 2017) and has already helped show when genes are essential (Hutchison et al. 2016), when gene overlap is non-essential (Jaschke et al. 2013) and, in general, that the “genomes encoding natural biological systems can be systematically redesigned and built anew in service of scientific understanding or human intention” (Chan et al. 2005). Practical applications of genome-scale reverse genetics approaches are also being pioneered, most notably in vaccine development (Coleman et al. 2008; Dormitzer et al. 2013; Desselberger 2017). Synthesis has also been used to reveal and better understand more subtle puzzles ranging from retrotransposon activity (Han and Boeke 2004) to whether the amount of conserved information encoded in a natural DNA sequence exceeds the known functions assigned to the sequence (Schneider and Stormo 1989).

Thus and taken together, we envisioned a more formal and systematic approach for determining if all ORFs in a given genome have been discovered. Rather than searching for individual phenotypes associated with each remaining potential cryptic ORF, we start by accepting a presumption that all cryptic ORFs have, individually and collectively, no functions whatsoever. Then, by using reverse genomics to systematically generate a derivative genome designed to test the starting presumption, we purposefully and efficiently identify if any further functions remain to be discovered or, if none are found, definitively declare that no such functions remain unknown. While the starting presumption may be widely accepted – otherwise any still-hidden genes would have been found already by established methods – the presumption itself seems never to have been tested by direct experiment. More importantly, if we can formally demonstrate that no further functional information is encoded in a natural genome, albeit with the simplest model genome as a first example, then we can demonstrate a method by which genetics, as a discovery science, can approach “completeness” in both an empirical and formal sense (Stent 1969).

## Results

### Identifying conserved ORFs in øX174

To directly test the completeness of the protein-coding ORF annotation of the bacteriophage øX174 genome we compared the ORFs of 23 *Bullavirinae* genomes to identify if any highly conserved ORFs could be detected. We focused on evolutionarily conserved ORFs because they are more likely to encode functions, such as protein-expression, compared to non-conserved ORFs (Carvunis et al. 2012).

We first filtered the set of all possible ORFs to include only those beginning with one of the three most common start codons in the *E. coli* host (ATG, GTG, and TTG) (Hecht et al. 2017;Firnberg et al. 2014) and preceded by an identifiable Shine-Dalgarno motif. Our analysis identified a set of 83 conserved ORFs shared between the øX174 genome and at least one other *Bullavirinae* genome (Fig. 1C). Eleven of these identified ORFs were the known protein-coding ORFs of the *Bullavirinae* subfamily (A, A*, B, C, D, E, F, G, H, J, K). We designated the remaining set of 72 ORFs as the cryptic ORFs of øX174 because they lack prior evidence of protein expression and function.

We ranked each of the 83 identified ORFs (11 known plus 72 cryptics) by the number of times each was detected by computational analysis of the 23 genomes studied, finding a range of conservation among the set (Fig. S1 and Fig. S2). Eight of the eleven known protein-coding ORFs clustered as the highest scoring due to their detection by all computational tools in a majority of the 23 searched genomes (Fig. S3). Known protein-coding ORFs A*, E, and K had much lower scores similar to those of eight cryptic ORFs (2, 13, 15, 29, 31, 46, 46, and 57) due to their detection by only some of the analysis methods (Fig. S3). We manually added 10 more cryptic ORFs that are known to be conserved across the *PhiX174microvirus* genus (Fig. S2) (Godson et al. 1978), leading to 82 candidate cryptic ORFs total.

### Design and characterization of a genome encoding only cryptic ORFs

We designed a variant øX174 genome in which the start codons initiating translation of the 11 known essential protein-coding ORFs are simultaneously disrupted. We called the resulting genome “cryptX174” as it should only express proteins from cryptic ORFs (Fig.1D). Our goals in making cryptX174 were to test the validity of the established start codons and to construct a template that contains all ORFs but for the 11 known protein-coding ORFs; we wanted a genome whose design better supports detection of peptides expressed at very low levels. Practically, the cryptX174 design changes the ATG start codons of protein-coding ORFs A, A*, B, C, D, E, F, G, H, J, K to either ATA or ACG (Table S2); specific alternate codons were selected to have the least impact on the amino acids coded in the other five reading frames.

We tested the viability of the cryptX174 design by *in vitro* construction followed by transfection into host *E. coli* C cells. We found that transfection of the cryptX174 genome did not produce observable plaques, as expected. CryptX174 non-viability suggests that a disruption in at least one of the essential protein-coding ORFs cannot be compensated for by an upstream or downstream in-frame start codon.

We next sought to determine if any cryptic ORFs are expressed. Given that the cryptX174 genome did not encode viable phage we used cell-free transcription and translation to measure protein production directly. We generated linear genomes from both the wild-type and cryptX174 templates, adding a terminal T7 promoter and terminator to each. We used mass spectrometry to measure cell-free protein production from both the wild-type and cryptX174 templates. We identified peptides from 8 of the 11 known protein-coding ORFs and cryptic ORF 75. Peptides from cryptic ORF 75 were produced from both wild-type øX174 and cryptX174 templates, and the N-terminal ORF 75 peptide (fMSIITPK) was observed unprocessed with a N-formylmethionine modification still intact (Table 1). N-formylmethionine is a marker of translation initiation in *E. coli* that is usually cleaved off quickly *in vivo* (Meinnel, Mechulam, and Blanquet 1993); the presence of a N-formylmethionine modification on the ORF 75 N-terminal methionine gave us confidence that this peptide arose from ORF 75-specific translation initiation and was not an internal peptide encoded by another ORF. N-formylmethionine peptides produced from genes A, D, and H were also observed, supporting the established start sites for these genes (Table 1).

**Table 1.**
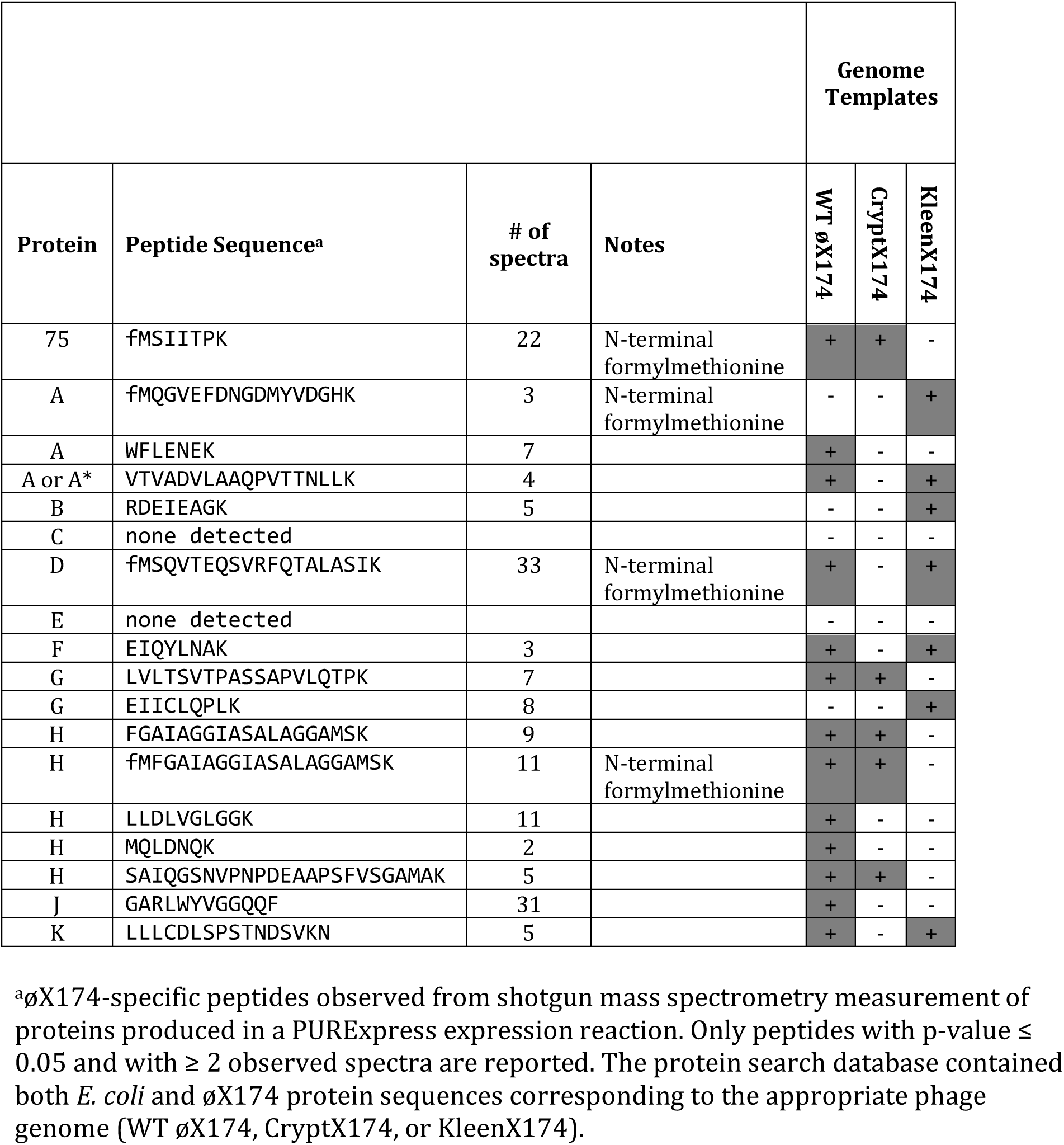
Peptides observed by mass spectrometry following cell-free expression from wild-type and synthetic genome templates

### Design and characterization of a genome “cleaned” of cryptic ORFs

We next designed a variant øX174 genome in which as many cryptic ORFs are simultaneously disrupted, as possible. To do so we introduced silent mutations that either: (1) changed the start codon to any codon but ATG, GTG, or TTG, or (2) weakened the putative ribosome binding site by reducing the frequency of A and G nucleotides, or (3) introduced a premature stop codon near the 5’-end of the ORF. In all cases we did not alter the protein coding sequence for any of the 11 known øX174 genes. By this approach we were able to design disruptions for 71 of the 82 cryptic ORFs, including ORF 75 (Table S1). We called the resulting genome “kleenX174” (Fig. 1E); the kleenX174 genome differs in only 120 positions from the wild type and has an unchanged GC-content (Fig. 2A).

**Figure 2.**
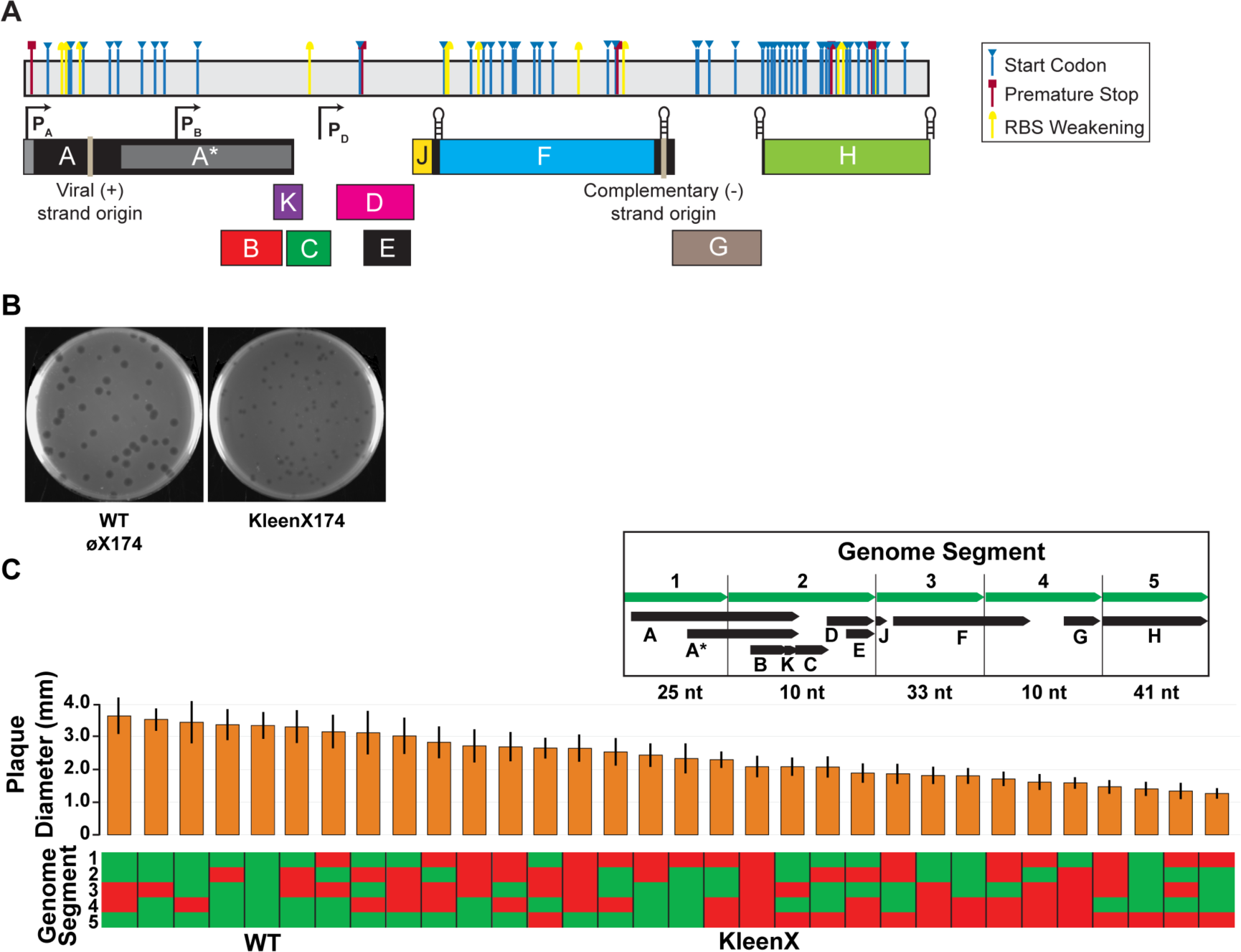
A “clean” øX174 genome devoid of most cryptic ORFs is viable but has reduced fitness. (A) linear depiction of kleenX174 genome showing the locations and modes of cryptic ORF disruption; see S1 Table for detailed information. (B) plaques of wild-type and kleenX174 phage. (C) plaque diameter of wild-type/kleenX174 chimeras, arranged from largest average plaque size to smallest. Vertical bars represent one standard deviation from n=50 plaque measurements. Each chimeric phage consists of five genome segments chosen from a mixture of wild-type genome segment (green) and kleenX174 modified genome segments (red). (Inset) The boundaries of the five genome segments, protein-coding ORFs found in each segment, and total number of nucleotide differences between wild-type øX174 and kleenX174 genome sequences in each segment.

As before, we constructed the kleenX174 genome by *in vitro* construction followed by transfection into host *E. coli* C cells, which in this case resulted in plaques (Fig. 2B). We found that kleenX174 plaques had a smaller diameter (2.1±0.3 mm) compared to wild-type plaques (3.3±0.4 mm). Smaller plaque sizes generally imply slower growth rates and lower fitness (Abedon and Yin 2009).

### KleenX174 chimera design and phenotypic measurements

To map the kleenX174 growth defect we made chimeric genomes by sectioning the øX174 genome into five segments and assembling 32 chimeras containing either the wild-type or kleenX174 sequence for each segment. We used plaque sizes for each chimeric phage to identify which of the 120 synonymous mutations within the kleenX174 genome design might contribute to a reduced plaque size phenotype. Measured diameters of the resulting plaques from each chimeric genome showed that segment five, encompassing all of gene H, was implicated as the loci within kleenX174 most responsible for its small-plaque phenotype (Fig. 2C).

### Identification of proteins produced from kleenX174

We next wanted to identify the proteins expressed from the kleenX174 genome. We used cell-free expression to produce proteins from linearized kleenX174 template and measured proteins via mass spectrometry. We identified peptides from five of the eleven known øX174 genes (Table 1). We did not detect ORF 75 peptides; the start codon of ORF 75 is disrupted in kleenX174 (ATG>ACG). We observed N-formylmethionine peptides produced from genes A and D (Table 1), further supporting the annotated start sites for these genes. We did not detect H protein peptides from kleenX174 template but did from both cryptX174 and wild-type templates. We hypothesized that the lack of detectable H protein peptides from kleenX174 template along with the low fitness of chimeric phage containing kleenX174 segment 5 might both be due to decreased protein H production (Fig. 2C).

### Evolutionary adaptation of kleenX174 reveals a functionally important codon

Refactored genomes can serve as a starting point for experimental adaptions that reveal design problems or novel functional elements (Springman et al. 2012). Thus, to better understand the kleenX174 growth defect we passaged the phage for 48 generations, selecting for faster growing mutants. Following this process, sequencing indicated that one change, kleenX174(2939C>T), was most abundant in both bulk culture and individually sequenced plaques (Fig. 3A). Moreover, we observed the 2939C>T mutation in repeated serial passage experiments. We isolated pure kleenX174(2939C>T) phage stock and measured the effect of the 2939C>T mutation on growth rates. We found that the growth rate and plaque size of the kleenX174(2939C>T) mutant was restored to that of wild type (Fig. 3B and C).

**Figure 3.**
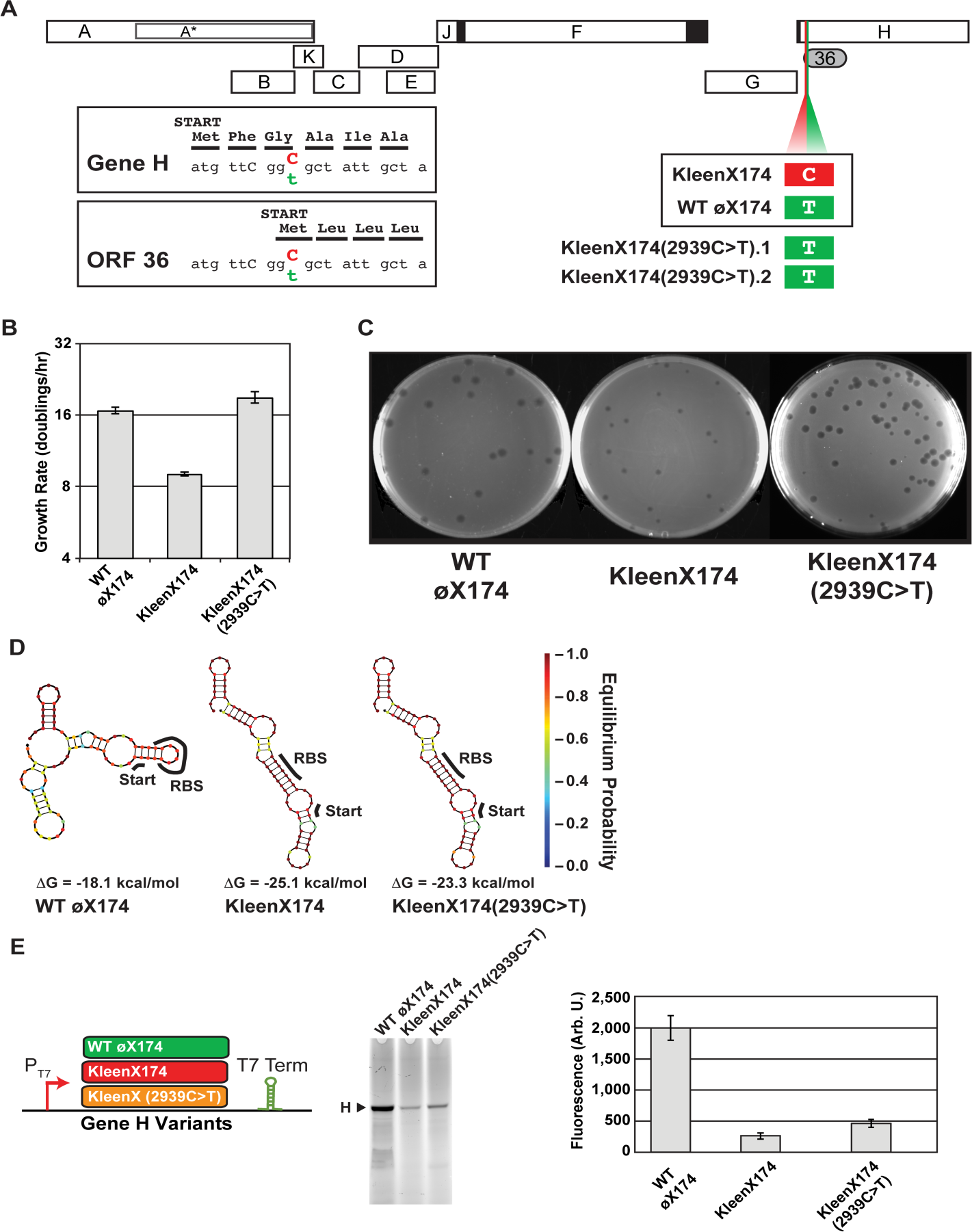
Evolutionary adaptation of kleenX174 to high-growth rate results in a reversion (2939C>T) and increased gene H expression. (A) the mutation 2939C>T observed in two independent adaptation experiments both restores the putative start codon of cryptic ORF 36 (GCG>GTG) and silently changes the third codon of gene H (GGC>GGT, Gly>Gly). (B) evolved KleenX174(2939C>T) grows faster than ancestral kleenX174 in liquid culture (population doublings per hour). (C) the kleenX174(2939C>T) mutation also recovers plaque size. (D) RNA structure predictions of gene H sequence variants from wild-type øX174, kleenX174 and mutant kleenX174(2939C>T). NUPACK lowest-energy RNA structures generated from an 83 nt window surrounding the gene H start codon. (E) Protein H production from kleenX174(2939C>T) is increased compared to ancestral kleenX174 genome. Protein H produced from synthetic dsDNA templates containing either wild-type øX174, kleenX174, or kleenX174(2939C>T) gene H sequence plus 20 bp upstream sequence identical to genome background, flanked by T7 promoter (P_T7_) and terminator sequences (T7 Term). PURExpress reactions run with 0.8 nM template were separated on SDS-PAGE followed by fluorescence detection of BODIPY-FL tagged lysine incorporated into proteins produced during the transcription/translation reaction. Error bars represent one standard deviation (n=3).

The 2939C>T mutation is located within segment 5 from the chimera experiment and appears to both revert the start codon of cryptic ORF 36 (from G<u>C</u>G to G<u>T</u>G) and, in a different reading frame, cause a synonymous glycine mutation (from GG<u>C</u> to GG<u>T</u>) in the third codon of gene H (Fig. 3A). GGC is used more frequently than GGT in *E. coli* (0.35 versus 0.27), suggesting this substitution might modulate gene H translation rate.

To explore the effect of the 2939C>T mutation on gene H mRNA structure we performed RNA folding simulations of the 5’-ends of gene H sequence variants. We found that the kleenX174 gene H structure is predicted to be -7 kcal/mol (28%) more stable than the wild type and has a different predicted base-pairing pattern (Fig. 3D, left and middle structures), while the kleenX174(2939C>T) mutant retains the altered RNA base-pairing structure of kleenX174 but should be less stable (Fig. 3D, right structure). We also found that the predicted gene H mRNA structural stability in the kleenX174 design is outside the range predicted using the gene H sequences from all 23 *Bullavirinae* genomes (Fig. S7). Higher mRNA stability around the start codon has been implicated in decreased protein expression (Goodman, Church, and Kosuri 2013).

To test the hypothesis that the small plaque phenotype of kleenX174 is due to decreased gene H protein production we created three synthetic versions of gene H matching those of wild-type øX174, kleenX174, and kleenX174(2939C>T). We measured H protein production from each template in a cell-free expression system (Fig. 3E). We found that H protein production from a kleenX174 template was only a fraction of wild-type levels (13%) and that the 2939C>T mutation almost doubles the amount of H protein produced (23%) (Fig. 3E).

## Discussion

We definitively determined by perturbation design, measurement, and evolutionary adaptation that the conventional annotation of functional genes in the øX174 genome is complete. Specifically, we found that 71 of 82 highly conserved cryptic ORFs within the øX174 genome are simultaneously dispensable for growth at wild-type rates and do not encode detectable amounts of protein. Even the five cryptic ORFs (2, 29, 31, 46, and 57) that have conservation scores greater than known protein-coding ORFs can be disrupted without apparent phenotypic effect (Fig. S1). We acknowledge that 11 of 82 conserved cryptic ORFs cannot be disrupted without also altering one or more known essential genes and were therefore not tested by ORF disruption directly. However, for each of these cryptic ORFs no peptides specific to their expression were detected by mass spectrometry. Disruption of one cryptic ORF (75) for which expressed peptides were detected by mass spectrometry had no observed impact on phenotype or fitness.

Evolutionary adaptation for faster-growing mutants derived from a synthetic genome devoid of most conserved cryptic ORFs revealed a consistent reversion that partially restored wild-type translation rates for a known essential gene (H). Gene H is a DNA pilot protein that is present at 12 copies per capsid (Fane et al. 2005) and needed for genome injection into host cells (Sun et al. 2014). The rarer glycine codon in the wild-type gene H N-termini could be important for slowing translation if needed to ensure correct folding, timing during infection (Plotkin and Kudla 2011), or level matching between B and H proteins during capsid assembly (Cherwa, Young, and Fane 2011).

We note that the so-selected reversion also restores a potential GTG start codon for cryptic ORF 36 (Fig. 3). The ORF 36 start codon and sequence is well conserved across 19 of the 23 genomes analyzed (Fig. S4 and Fig. S5). However, despite the presence of an upstream RBS, strong start codon, and strong family conservation, we did not observe any peptides from ORF 36 in our experiments and conclude that ORF 36 does not produce protein.

From a methods perspective, we note that many of the proteins produced via the cell-free expression system had intact N-terminal formylmethionine residues (Table 1). The PURE transcription and translation system used in this work appears to lack the peptide deformylase activity necessary to remove N-terminal formyl groups (Shimizu et al. 2001). Such residues serve as hallmarks of translation initiation in bacteria but are typically quickly removed *in vivo*, thereby masking true translation initiation sites (Laursen et al. 2005). Established methods such as ribosome footprinting (Ingolia et al. 2014) identify translation initiation sites by isolating ribosomes in the vicinity of mRNA start codons. However, detection of formylmethionine on peptide N-termini provides direct biochemical evidence of translation initiation. The PURE system therefore shows significant promise as a tool for systematically mapping translation initiation, especially compared to methods that require peptide enrichment (McDonald and Beynon 2006) or the use of knockout strains and translation inhibitors (Spector et al. 2003).

Even given the increasing pace of progress in basic and applied biological sciences over the last 150 years, the scope and complexity of the natural living world still feels infinite. However, we note that genomes encoding natural living systems are finite in length; the rate at which novel functional genetic information is “written” into genomes is limited by how quickly mutations can be selected for and fixed within a competing population; and, the pace at which encoded functional genetic information is “lost” from genomes is greater than zero, as determined by spontaneous mutation rates and functional selection frequencies. Taken together, these three facts – (i) an information storage system of finite capacity, (ii) a finite rate of information encoding, and (iii) a non-zero rate of information loss – practically bound the amount of functional information that can be encoded in any natural genome at any point in time. Thus, the science of discovering and understanding functional genetic information encoded in natural living systems (i.e., genetics) should itself be formally finite and bounded. Herein, we have explored such a finite framing by starting from the presumption that the existing discovery science for one natural genome had been completed. Although the understanding for almost all other natural living systems is now very far from complete, we submit for consideration that the underlying framing should nevertheless hold, and suggest that the science of genetics may progress to completion more quickly by more systematically complementing classical forward discovery-based approaches with reverse approaches that definitively demonstrate what does not matter. For engineers or others interested in refining and repurposing natural genetic sequences we note that the methods applied here can collectively serve to validate DNA by demonstrating definitively what molecules are encoded in any given DNA sequence.

## Methods

### Bacterial strains, growth conditions, and plaque assays

The bacterial strain *E. coli* C (ATCC 13706) grown at 37C in phage LB was used throughout (Rokyta, Abdo, and Wichman 2009). Transformed *E. coli* C cells were regrown for 1.5 hours in phage LB followed by mixing 10 µL transformed cell serial dilutions with 200 µL untransformed *E. coli* C grown overnight to saturation. The mixed cells were plated using the double-layer agar plate method (15 mL bottom-layer 1.2% agar, 5 mL top-layer 0.7% agar) with either phage LB or TK agar plates (Fane and Hayashi 1991; Jaschke et al. 2012). Plates were incubated at 37C for 16-18 hours before visualization. Plaque sizes were measured using Adobe Photoshop CS6 Analysis->Measurement command from images of plates containing between 40-100 plaques.

### ORF prediction

Four established gene prediction tools were used in combination to find ORFs in the øX174 genome: GLIMMER (Delcher et al. 2007), GeneMark (Besemer and Borodovsky 2005) using the GeneMark.hmm PROKARYOTIC (version 2.10b) algorithm, EasyGene (Larsen and Krogh 2003), and Prodigal (Hyatt et al. 2010). We analysed genomes from 23 different *Bullavirinae* virus subfamily phages from three different genera: the *PhiX174microvirus* phages: WA11 (DQ079895.1), WA10 (DQ079894.1), WA4 (DQ079893.1), S13 (M14428.1), øX174 (NC_001422.1), NC56 (DQ079892.1), NC51 (DQ079891.1), NC41 (DQ079890.1), NC37 (DQ079889.1), NC16 (DQ079888.1), NC11 (DQ079887.1), NC7 (DQ079886.1), NC5 (DQ079885.1), NC1 (DQ079884.1), ID45 (DQ079883.1), ID22 (DQ079881.1), and ID1 (DQ079880.1); the *G4microviruses*: ID18 (NC_007856.1), G4 (NC_001420.1), and ID2 (NC_007817.1); and the *alpha3microviruses*: alpha3 (NC_001330.1), St-1 (NC_012868.1), and WA13 (NC_007821.1). These genome sequences were analyzed for ORFs initiating from the strongest three start codons in *E. coli*: ATG, GTG, or TTG. In addition to computationally identified ORFs we also identified ten ORFs by reference to past literature that manually predicted potential protein-producing ORFs in the øX174 genome based on RBS location upstream of ATG start codons (Godson et al. 1978).

### Phage genome design and construction

All known commercial øX174 preparations have sequences that differ from the canonical Sanger øX174 sequence used as the basis for our kleenX174 design (Smith et al. 2003). Thus, we built from scratch the wild-type øX174 genome corresponding to the original Sanger 1977 sequence, as defined in GenBank (Sanger et al. 1977). Specifically, we downloaded the NC_001422.1 sequence, split it into five approximately equal size pieces with 60 bp overhangs, synthesized, assembled using *in vitro* homologous recombination (Gibson et al. 2009), and transformed directly into the *E. coli* C host strain (Warren, 2011;Chung, Niemela, and Miller 1989) to recover viable phage.

To design the kleenX174 genome (GenBank: MF426914.1) the wild-type øX174 sequence was modified so that 71 of 82 predicted cryptic ORFs would be disrupted (Table S1). We constrained genome modifications to only incorporate silent changes, whereby the amino acid sequence of any overlapping known øX174 protein-coding ORF remained the same. We first attempted to modify the start codon of each cryptic ORF as we reasoned doing so would give the highest probability of successfully disrupting ORFs while minimizing the number of nucleotides altered. However, if a start codon could not be so-altered we instead changed the RBS of the gene to reduce A and G base frequency, or modified in-frame bases to create a premature stop codon as close to the start codon as possible. In 11 cases no silent changes could be made without also disrupting a known ORF and so these cryptic ORFs were left unchanged; such cases were concentrated in the area of the genome containing the overlapping essential genes A, A*, B, and K.

The kleenX174 genome was synthesized in five parts with 60 bp overhangs, assembled using *in vitro* homologous recombination (Gibson et al. 2009), and transformed directly into the *E. coli* C host strain (Warren, 2011;Chung, Niemela, and Miller 1989).

To design and build the cryptX174 genome (GenBank: MF426915.1) we analyzed the start codons of all known øX174 protein-coding ORFs to determine nucleotide changes that could be made to mutate each to another codon that was not ATG while minimizing the impact on all overlapping upstream and downstream codons in the five remaining reading frames. The so-specified cryptX174 genome was synthesized in three parts with 60 bp overhangs, assembled using *in vitro* homologous recombination followed by transformation directly into *E. coli* C cells.

### Chimeric genome design and construction

The Sanger 1977 wild-type øX174 and kleenX174 genomes were computationally split into five approximately equal sized segments with joints between segments placed in locations that did not have any modifications from the wild-type sequence. The areas around the synthetic DNA joints had this requirement so that they could be swapped seamlessly with the corresponding area of the wild-type genome during construction of chimeric genomes. Assembly of the 32 chimeric genomes was planned using the Teselagen web-application (Hillson, Rosengarten, and Keasling 2012). Five wild-type and five kleenX174 parts were amplified using primers specified by Teselagen (Table S3) and assembled from 30 fmol of each of the five pieces using *in vitro* homologous recombination. Assembled genomes were transformed directly into chemically competent *E. coli* C.

### Creating linear genome templates for cell-free expression and mass spectrometry

We linearized the genomes just following the stop codon of gene H and added 88 bp of the H/A-intergenic region and start of gene A to the 3’ end of each template. These modifications ensured that each linear genome contained uninterrupted sequences for all predicted ORFs (Fig. S2). We added 8.6 nM of each genome and 1U/µL Murine RNase inhibitor to the PURExpress expression system (New England Biolabs). Each genome was ran in a separate reaction using either ^12^C/^14^N-, ^13^C/^14^N-, or ^13^C/^15^N-lysine (Thermo Scientific Pierce), 25C for 16 hrs. All three reactions were pooled and digested with Lys-C, cleaned up, and ran on nanoLC-2D and LTQ-Orbitrap Velos (ThermoFisher) followed by analysis with Byonic v2.0-25 software. The peptide search database contained all *E. coli* protein sequences pooled with the 11 known øX174 proteins and the 82 cryptic ORFs identified in this work.

### Evolutionary selection of kleenX174 phage

To subject the kleenX174 phage to evolutionary selection for increased growth rate we followed an established adaptation protocol (Rokyta, Abdo, and Wichman 2009). Briefly, *E. coli* C was grown overnight at 37C in phage LB then split 1/100 into 25 mL of phage LB and regrown to A600=0.5, followed by the addition of 10^4^ KleenX174 phage (MOI = 2e^-5^). The infected culture was shaken for 40 minutes (~3 generations) and then chloroform was added to a final concentration of 4% (v/v). The culture was then centrifuged at 15,000xg for 15 minutes and the supernatant was removed to another tube, titred to determine phage concentration, and used to infect a fresh batch of *E. coli* cells at A600=0.5. After 16 passages (~48 generations) the lysate was used as template in rolling circle amplification (RCA) reactions and Sanger sequenced using a set of primers to cover the entire øX174 genome (Table S3). The lysate was also plated and isolated plaques were extracted into water and used as template for RCA followed by Sanger sequencing.

### Protein H production from synthetic DNA templates

Double stranded synthetic DNA templates corresponding to the wild-type øX174, kleenX174, and kleenX174(2939C>T) sequences were designed and synthesized (IDT) with added 5'T7 promoter and 3'T7 terminators, as recommended by NEB in the PURExpress kit. Cell-free *in vitro* expression reactions were run with 0.8 nM template at 37C for two hours under standard conditions with 1U/µL Murine RNase inhibitor (NEB) and FluoroTect BODIPY-FL labeled Lysine (Promega). Cell-free reactions were processed with RNase A (0.1 mg/mL) to remove unreacted tRNA, then run on 12% Bis-Tris SDS-PAGE using MES buffer (Life Technologies). Fluorescence was visualized using a Typhoon 9410 (GE) scanner with laser settings of Blue2 488 nm BP 520 normal sensitivity. Band volumes were calculated using ImageJ v1.49 (Schneider, Rasband, and Eliceiri 2012).

### mRNA structure predictions

RNA folding simulations were performed on an 83 nt window around the start codon of each analyzed ORF using the NUPACK web-server with default settings (Zadeh et al. 2011). Lowest-energy structures were reported. MUSCLE (Edgar 2004) multiple sequence alignments were also performed using the EMBL-EBI webservers <u>(http://www.ebi.ac.uk/Tools/msa/muscle/).</u>

## Data availability

All data generated or analyzed during this study are available from the corresponding authors on reasonable request. Sequences of kleenX174 (MF426914.1) and cryptX174 (MF426915.1) are available via NCBI Genbank.

## Acknowledgements

The authors acknowledge Jerome Bonnet, Pakpoom Subsoontorn, Monica Ortiz, Anwar Sunna, Ariel Hecht, Tom Williams, Heinrich Kroukamp, Jeff Glasgow, Marc Salit, Arend Sidow, and Joe Jacobson for helpful discussions and feedback. We acknowledge Ryan Leib and Chris Adams at the Vincent Coates Foundation Mass Spectrometry Laboratory, Stanford University Mass Spectrometry (http://mass-spec.stanford.edu) for assistance in protein analysis. This work was supported by students enrolled in the Stanford University Bioengineering Department Research Experience for Undergraduates (REU) program and the San Mateo High School Biotechnology Career Pathway Internship Program. PRJ was supported by a Canadian Natural Sciences and Engineering Research Council (NSERC) Postdoctoral Fellowship (PDF - 388725 - 2010) and Macquarie University’s Molecular Sciences Department, Faculty of Science, and Deputy Vice Chancellor (Research). Additional support was provided by the Stanford/NIST Joint Initiative for Metrology in Biology (jimb.stanford.edu) and a gift from Agilent Technologies, Inc.

## Author Contributions

DE and PRJ conceived the study. PRJ, GAD, KH, and DL performed experiments, analyzed data, and created the figures and tables. PRJ and DE wrote the manuscript. All authors read and approved the final manuscript.

## Supplementary Information

Supplementary Information.PDF

Table S1– PhiX174 ORF characteristics and modifications to generate kleenX174 genome design.xlsx

Data File S1 - P(T7)+WT_PhiX174_Gene_H+T7_min.gb

Data File S2 - P(T7)+KleenX174_Gene_H+T7_min.gb

Data File S3 - P(T7)+KleenX174(2939C_T)_GeneH.gb

## SUPPLEMENTARY INFORMATION

### CONTENTS

Supplementary Figures

Fig. S1 Computational ORF predictions.

Fig. S2 Computationally identified ORF locations on øX174 genome.

Fig. S3 Computational tool contributions to ORF prediction scores.

Fig. S4 Multiple sequence alignment of Gene H/ORF 36 regions from 23 *Bullavirinae* genomes.

Fig. S5 Multiple sequence alignment of ORF 36 from 19 *Bullavirinae* genomes.

Fig. S6 Simulated RNA folding structures of all known øX174 protein-coding ORFs in WT and KleenX174 genomes.

Fig. S7 Predicted gene H RNA structure in 23 *Bullavirinae* genomes shows KleenX174 and KleenX174(2939C>T) outside the normal range.

Supplementary Tables

Table S1 - PhiX174 ORF characteristics and modifications to generate KleenX174 genome design.xlsx

Table S2. Changes made to wild-type øX174 genome to produce CryptX174 design.

S3 Table. Oligonucleotides used in this work.

Supplementary Data Files

File S1. Wild type sequence gene H synthetic template for cell-free protein expression. Genbank file.

File S2. KleenX174 sequence gene H synthetic template for cell-free protein expression. Genbank file.

File S3. KleenX174(2939C>T) sequence gene H synthetic template for cell-free protein expression. Genbank file.

---

**S1 Fig.**
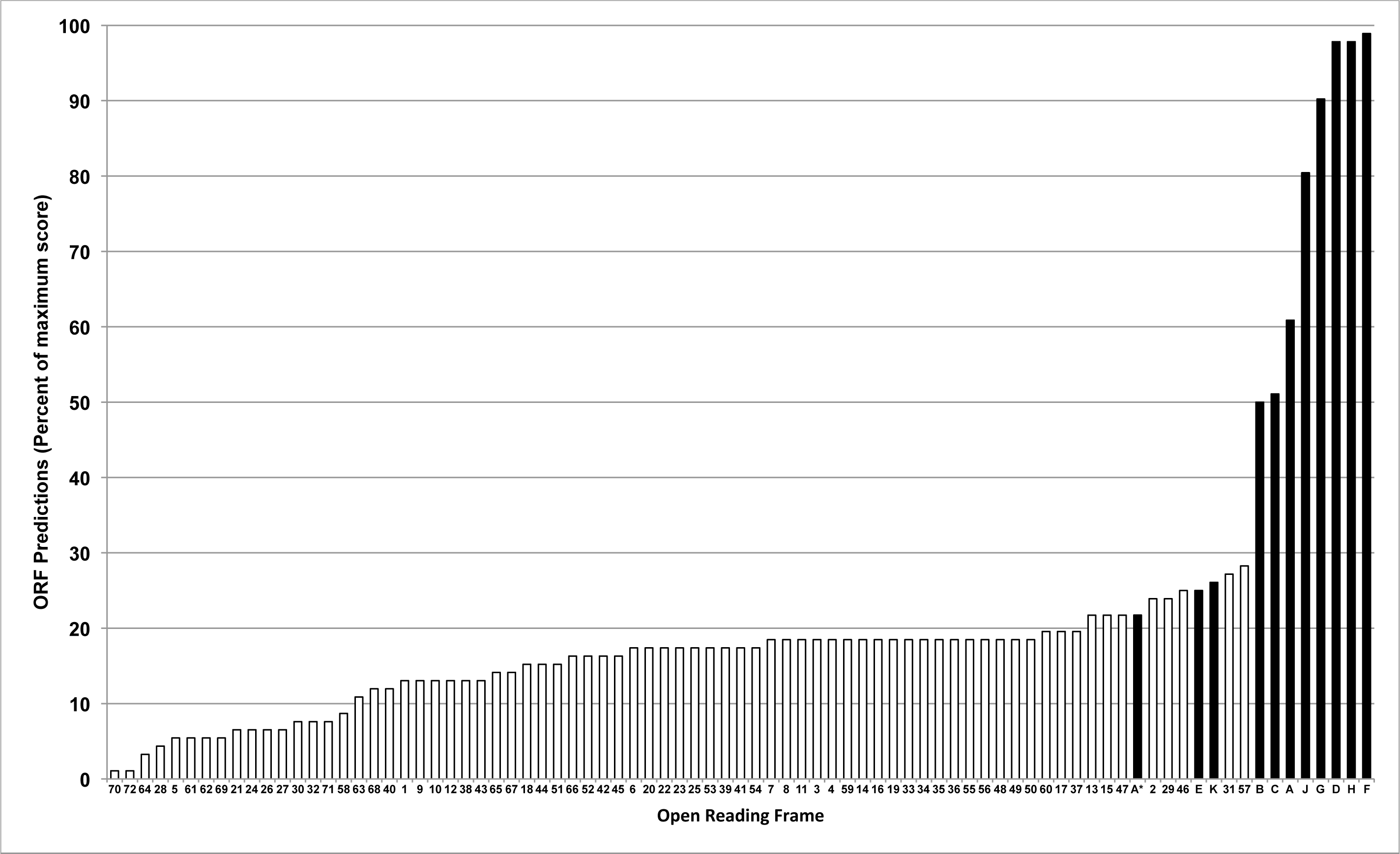
Computational ORF predictions. ORFs were given a score based on the number of times it was identified across 23 *Bullavirinae* genomes (92 possible identifications for each ORF, based on four computational tools and 23 genomes). Seventy-two cryptic ORFs (white bars) and 11 previously discovered øX174 protein-coding ORFs (black bars).

**S2 Fig.**
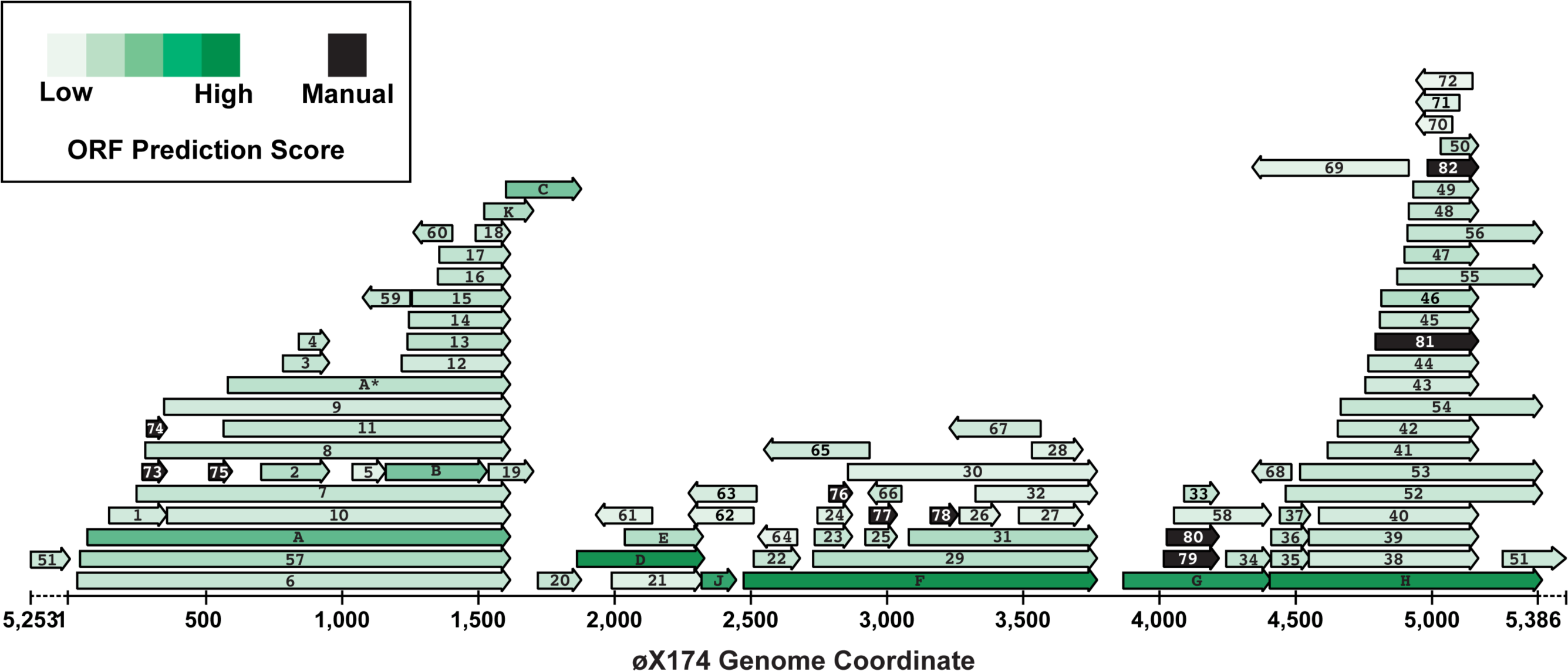
Computationally identified ORF locations on øX174 genome. ORFs were given a score based on the number of times it was identified across 23 *Bullavirinae* genomes (92 possible identifications for each ORF, based on four computational tools and 23 genomes). Black ORFs indicate 10 expert-curated cryptic ORFs [16].

**S3 Fig.**
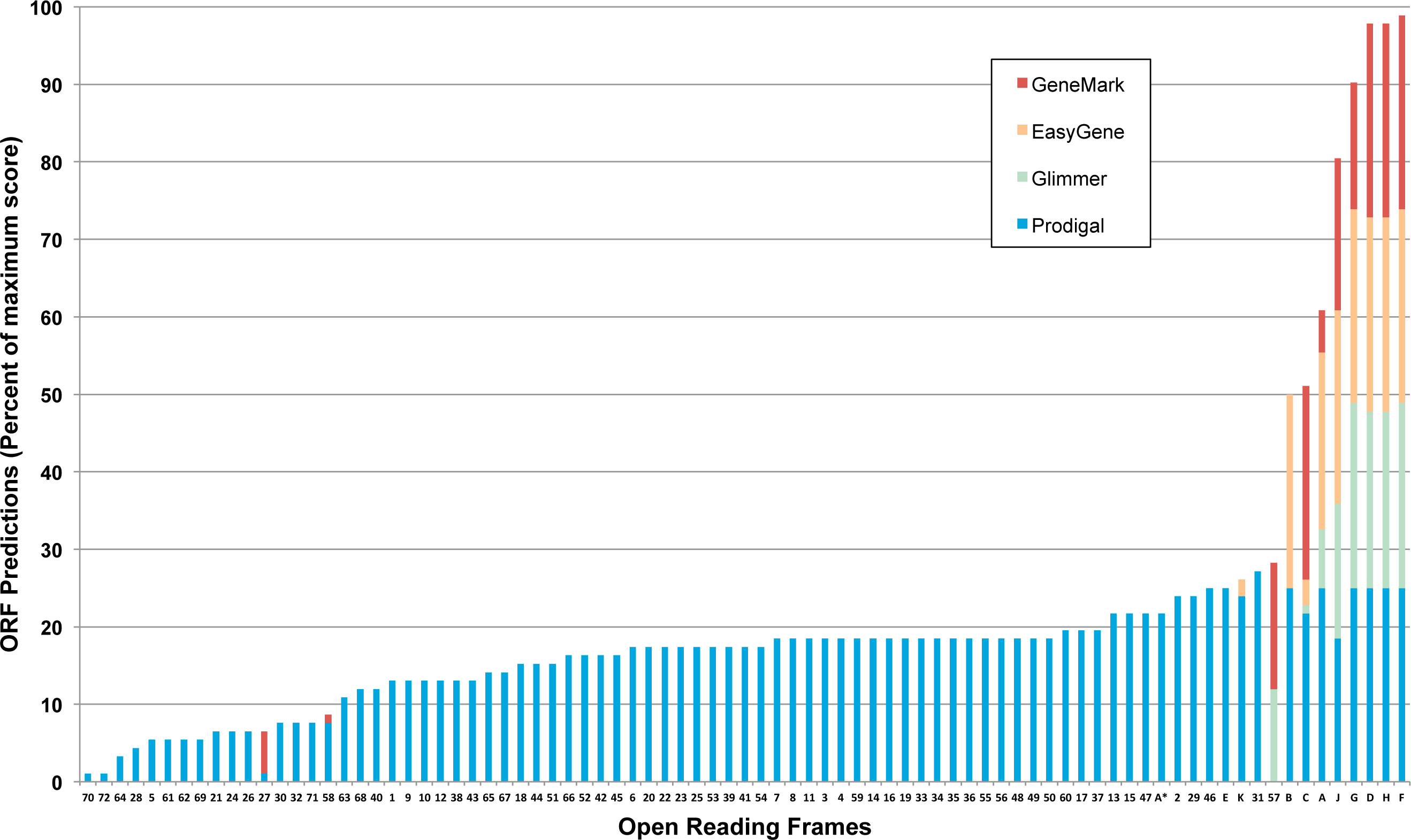
Computational tool contributions to ORF prediction scores. Identified ORFs were given a score based on the number of times it was identified across 23 *Bullavirinae* genomes (92 possible identifications for each ORF, based on four computational tools and 23 genomes). Four standard gene prediction tools were used: GLIMMER [38], GeneMark [39] using the GeneMark.hmm PROKARYOTIC (Version 2.10b) algorithm, EasyGene [40], and Prodigal [41].

**S4 Fig.**
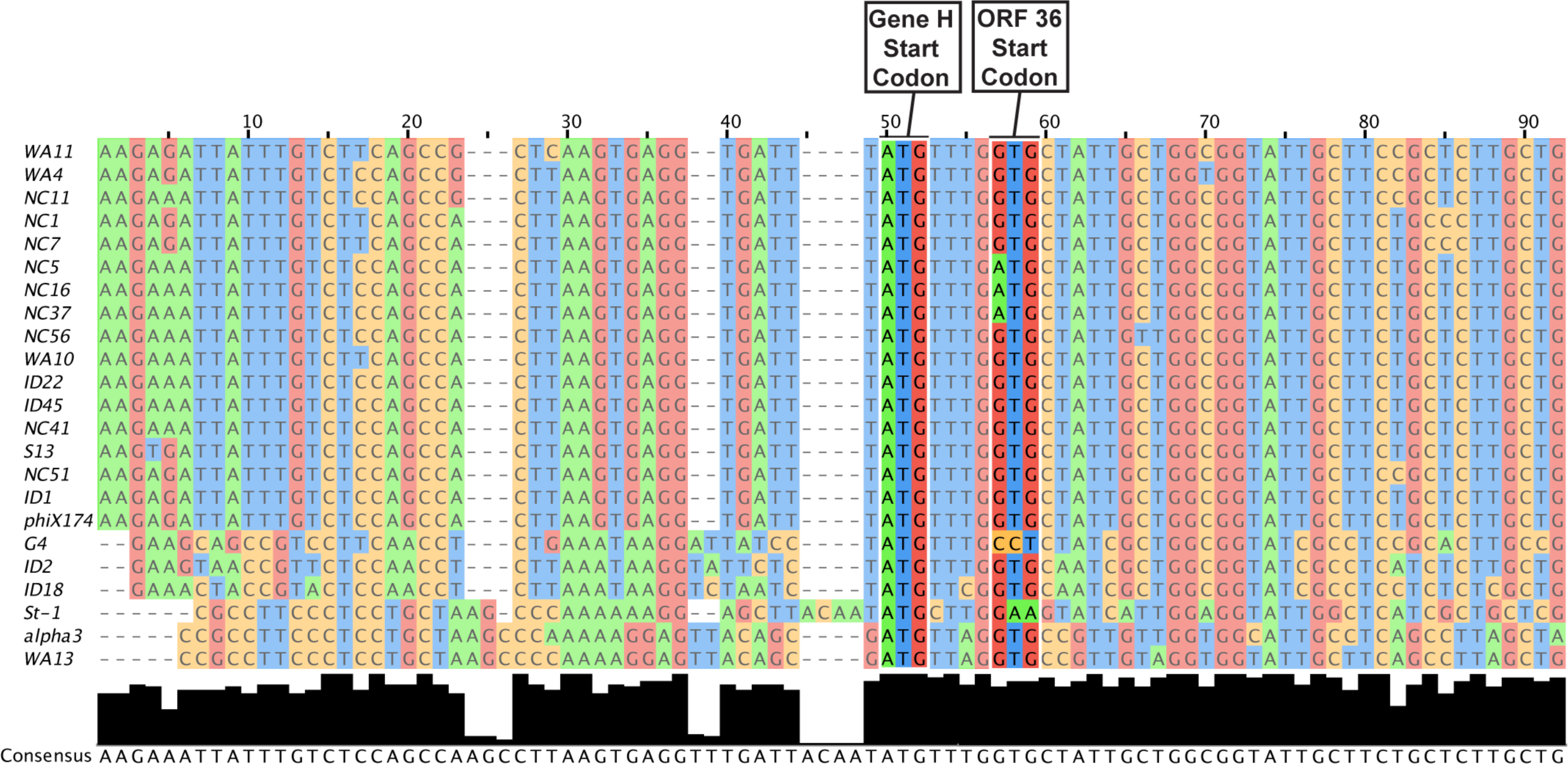
Multiple sequence alignment of Gene H/ORF 36 regions from 23 *Bullavirinae* genomes. Multiple sequence alignment of 83 nt centered on the start codon of gene H performed with MUSCLE multiple sequence alignment algorithm. Height of the black bars below each nucleotide represents degree of conservation within that column.

**S5 Fig.**
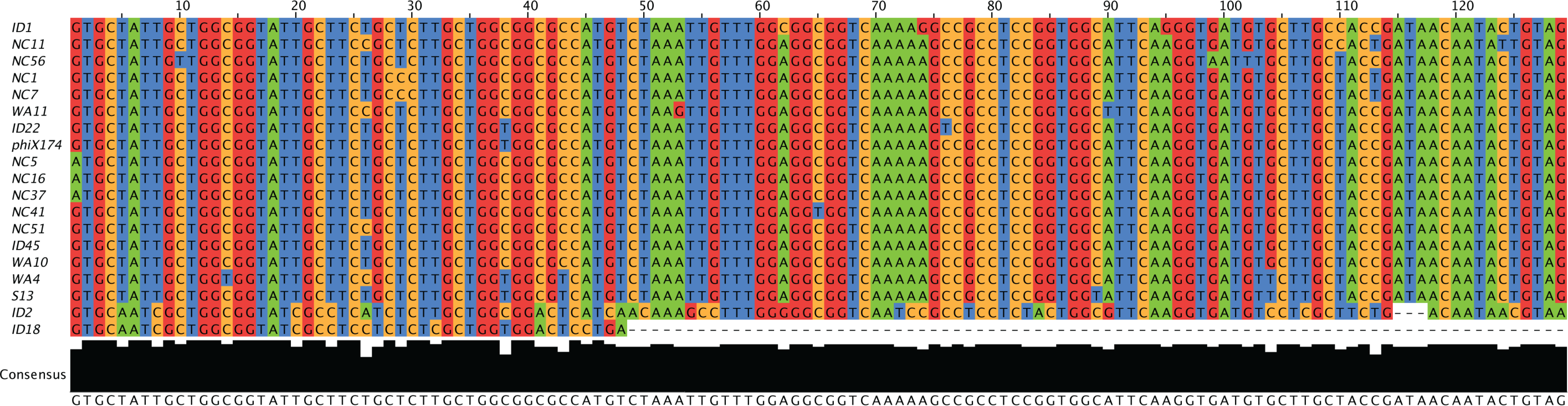
Multiple sequence alignment of ORF 36 from 19 *Bullavirinae* genomes. Multiple sequence alignment of ORF 36 performed with MUSCLE multiple sequence alignment algorithm. WA13 and alpha3 ORF36 sequences not included in alignment because their short length disrupted the alignment. G4 and st-1 ORF36 sequences lack strong start codons and were not included in the alignment. Height of the black bars below each nucleotide represents degree of conservation within that column.

**S6 Fig.**
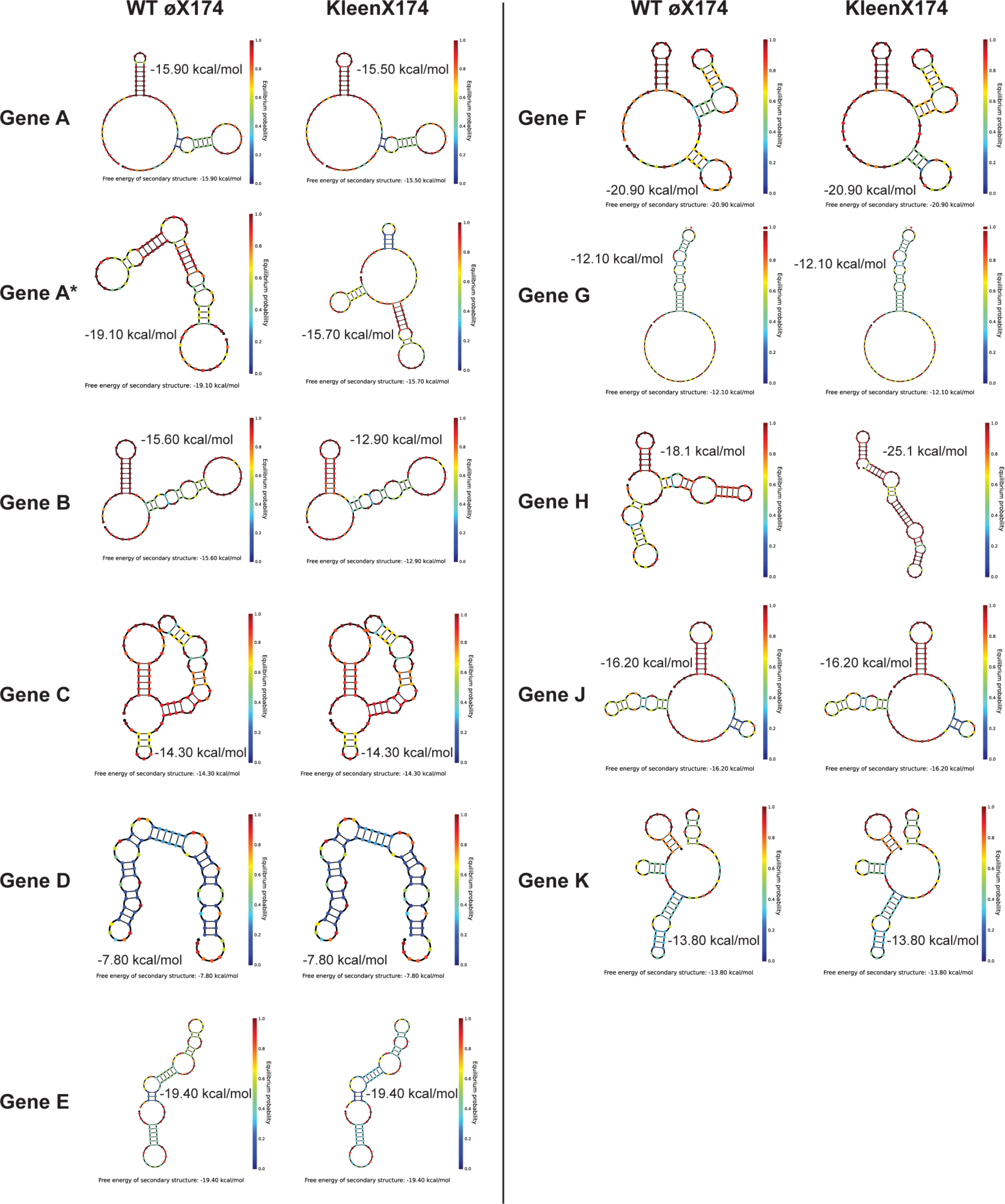
Simulated RNA folding structures of all known øX174 protein-coding ORFs in WT and KleenX174 genomes. NUPACK lowest energy RNA structure from 83 nt window surrounding each known øX174 gene using sequence variants from WT øX174 and KleenX174 genome sequences.

**S7 Fig.**
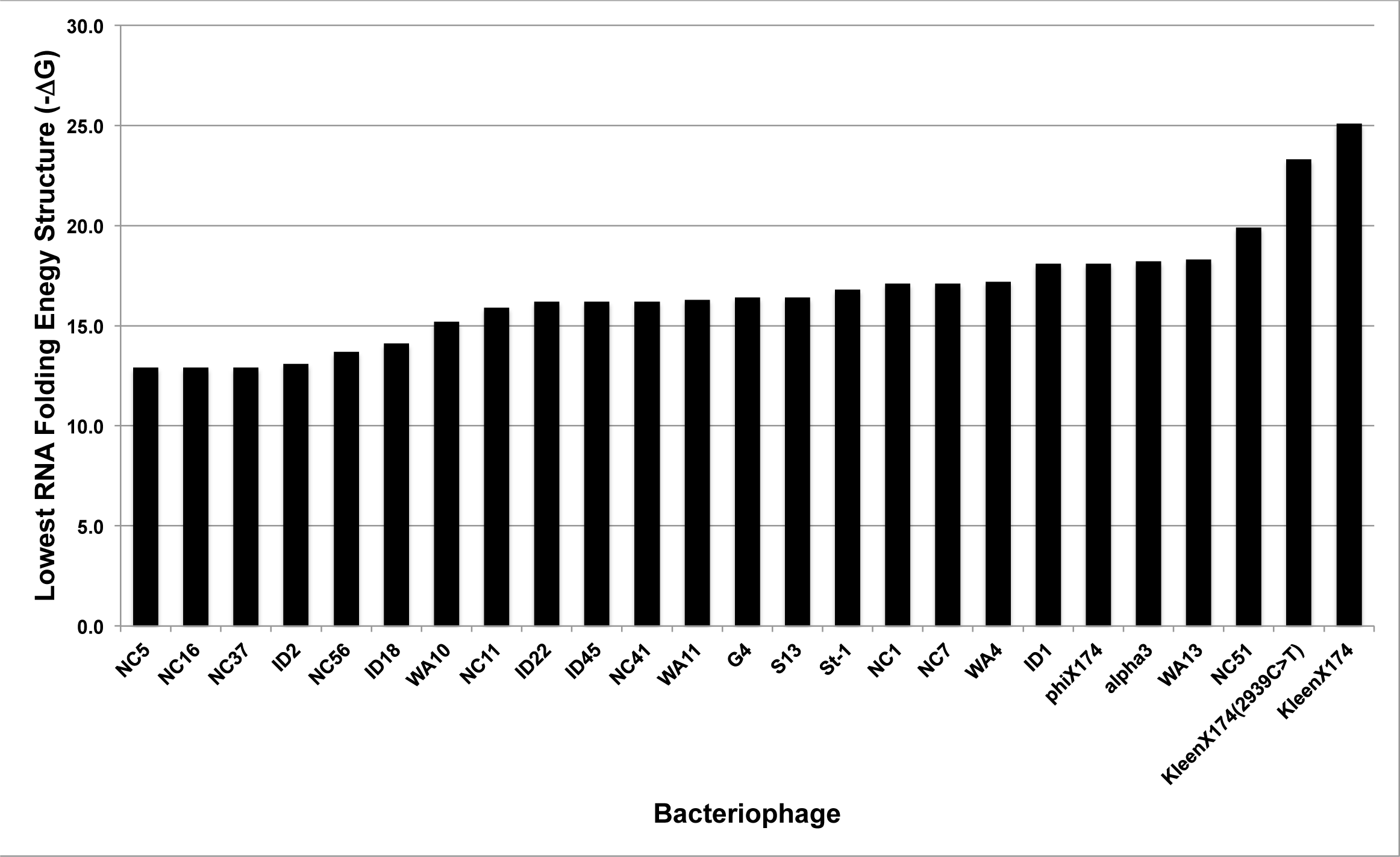
Predicted gene H RNA structure in 23 *Bullavirinae* genomes shows KleenX174 and KleenX174(2939C>T) outside the normal range. Lowest energy structures of RNA folding simulation performed with NUPACK using default parameters. Folding window of 83 nt centered on gene H initiation codon.

**Table S2.** Changes made to wild-type øX174 genome to produce CryptX174 design.

| Genome Coordinate<sup>a</sup> | WT Nucleotide | CryptX Nucleotide | Gene 1 | Codon Affected<sup>b</sup> | Amino Acid Change | Gene 2 | Codon Affected<sup>b</sup> | Amino Acid Change | Gene 3 | Codon Affected<sup>b</sup> | Amino Acid Change |
| --- | --- | --- | --- | --- | --- | --- | --- | --- | --- | --- | --- |
| 3982 | T | C | A | atg > aCg | Start (Met) > Thr | - | - | - | - | - | - |
| 4499 | G | A | A/A* | atg > atA | Start (Met) > Ile | - | - | - | - | - | - |
| 5076 | T | C | B | atg > aCg | Start (Met) > Thr | A/A* | tgg > Cgg | Trp > Arg | - | - | - |
| 53 | G | A | K | atg > atA | Start (Met) > Ile | A/A* | gag > Aag | Glu > Lys | - | - | - |
| 135 | G | A | C | atg > atA | Start (Met) > Ile | K | gag > Aag | Glu > Lys | A/A* | tga > tAa | Stop > Stop |
| 392 | G | A | D | atg > atA | Start (Met) > Ile | C | tga > tAa | Stop > Stop | - | - | - |
| 569 | T | C | E | atg > aCg | Start (Met) > Thr | D | tag > taC | Tyr > Tyr | - | - | - |
| 850 | G | A | J | atg > atA | Start (Met) > Ile | - | - | - | - | - | - |
| 1003 | G | A | F | atg > atA | Start (Met) > Ile | - | - | - | - | - | - |
| 2397 | G | A | G | atg > atA | Start (Met) > Ile | - | - | - | - | - | - |
| 2933 | G | A | H | atg > atA | Start (Met) > Ile | - | - | - | - | - | - |

<sup>a</sup>Coordinate in original Sanger øX174 genome sequence (Genbank NC\_001422.1)
<sup>b</sup>Capital letters represent mutated base positions in CryptX174 design.

**Table S3.**
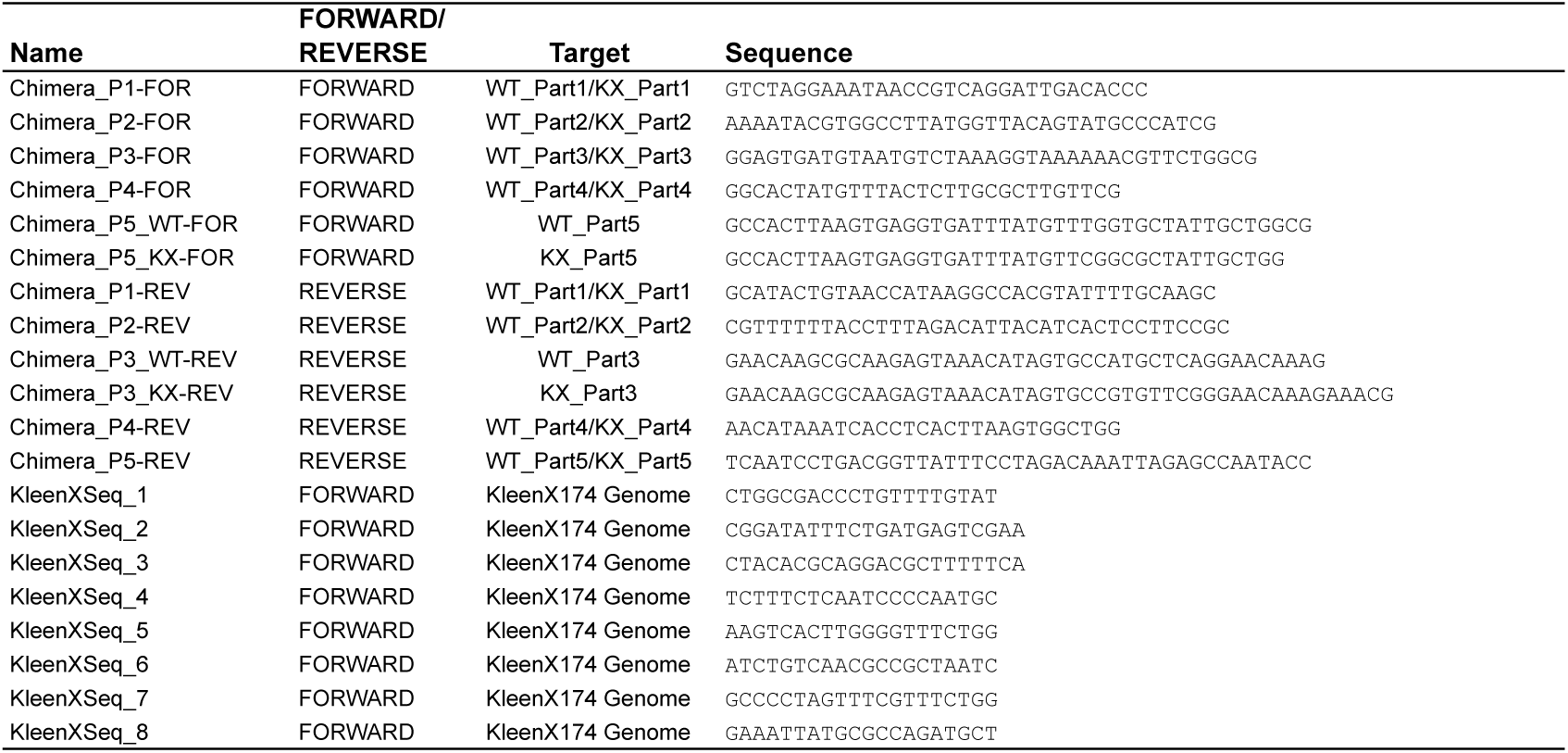
Oligonucleotides used in this work.

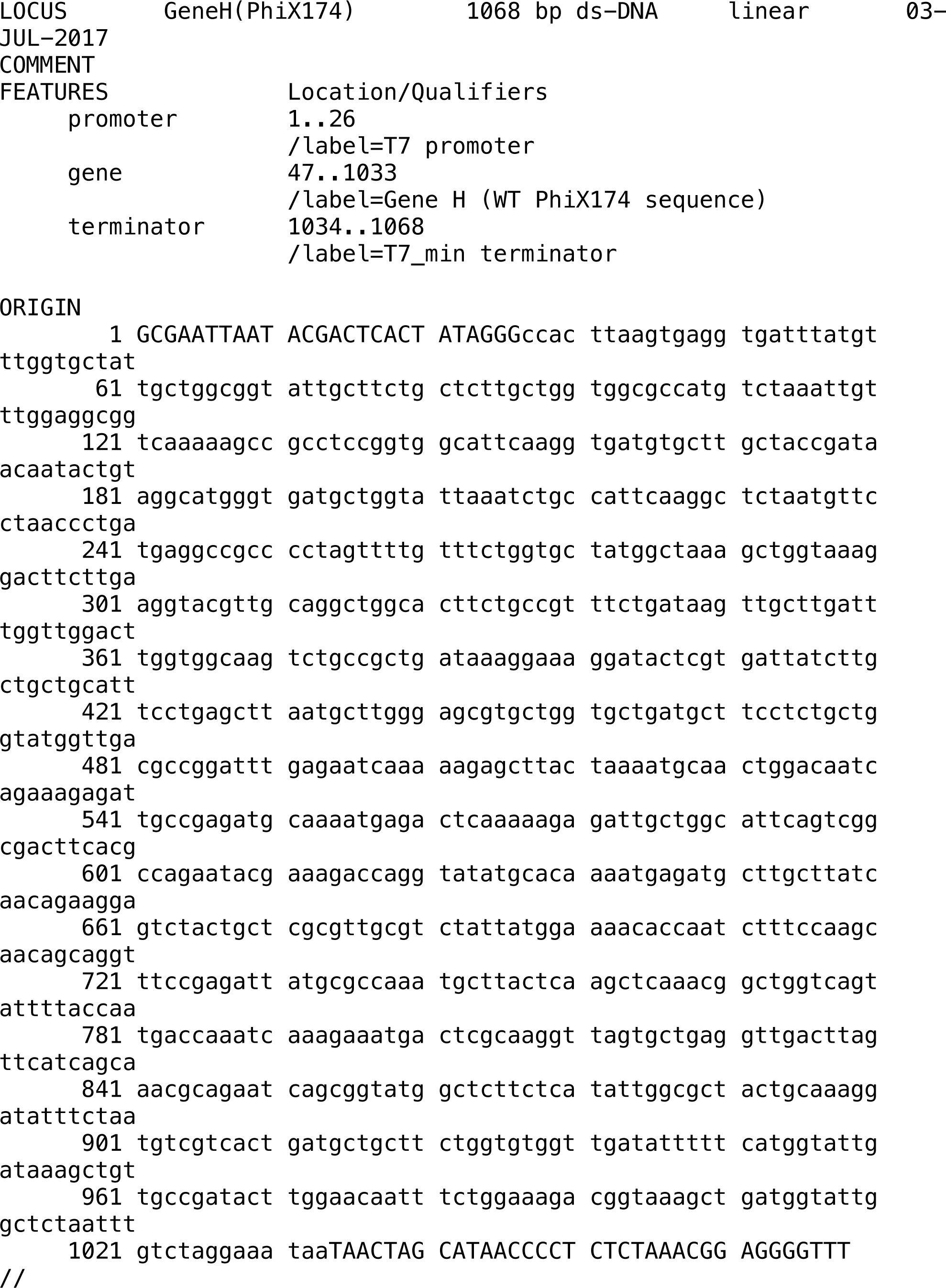

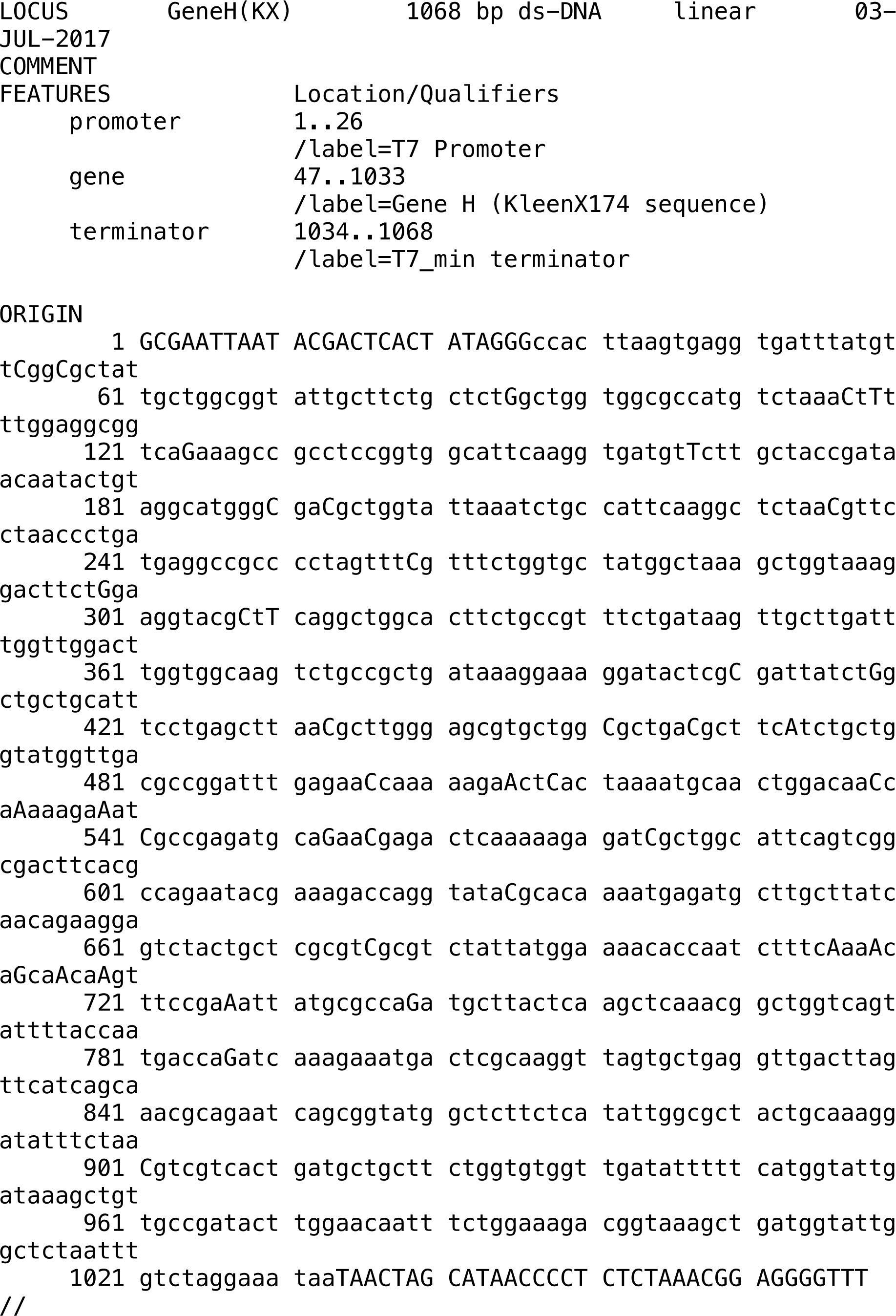

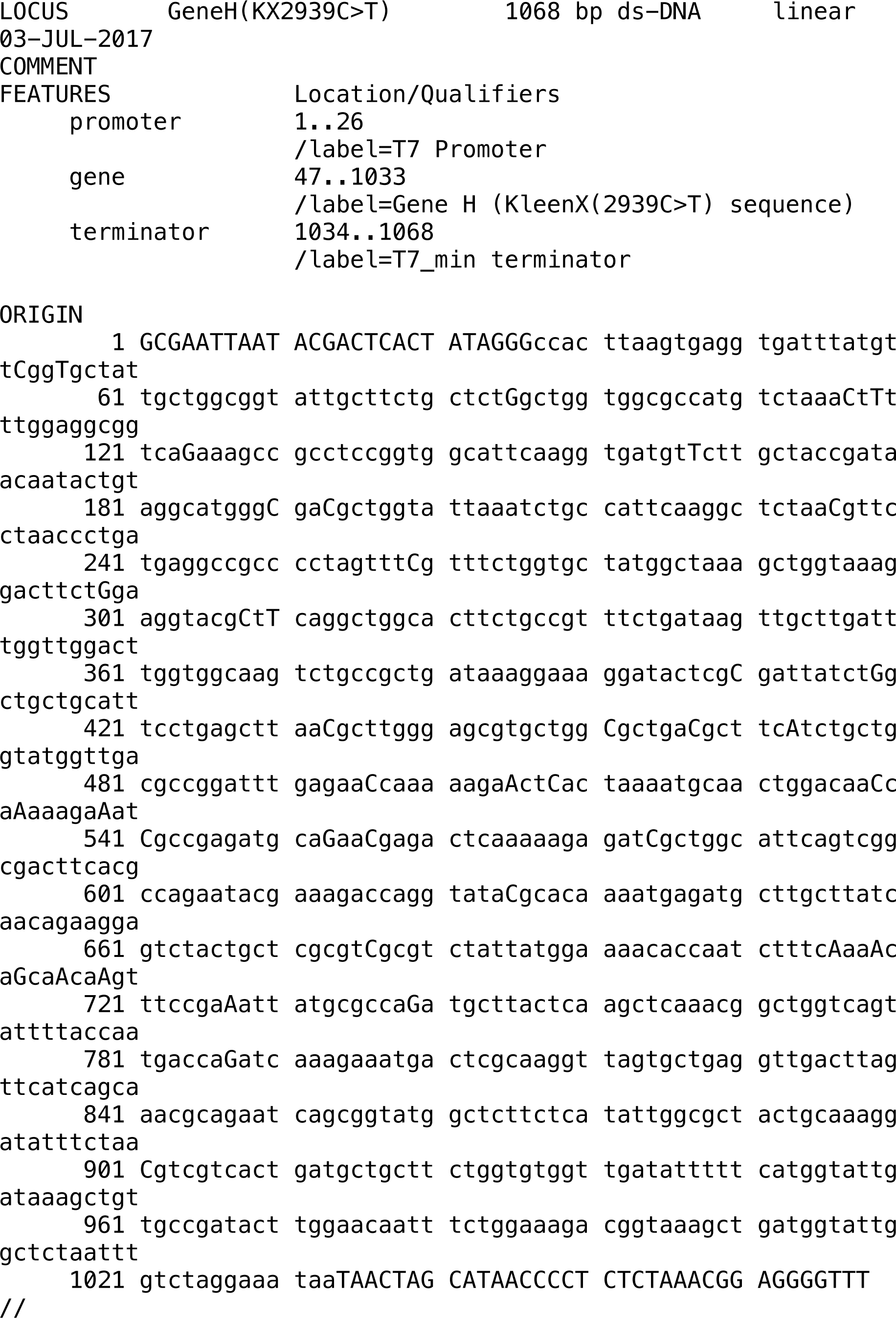

